# Assessing Per- and Polyfluoroalkyl Substances (PFAS) in Fish Fillet Using Non-Targeted Analyses

**DOI:** 10.1101/2023.09.01.555938

**Authors:** Anna K. Boatman, Jessie R. Chappel, Madison E. Polera, James N. Dodds, Scott M. Belcher, Erin S. Baker

## Abstract

Per- and polyfluoroalkyl substances (PFAS) are a class of thousands of man-made chemicals that are persistent and highly stable in the environment. Fish consumption has been identified as a key route of PFAS exposure for humans. However, routine fish monitoring targets only a handful of PFAS, and non-targeted analyses have largely only evaluated fish from heavily PFAS-impacted waters. Here, we evaluated PFAS in fish fillets from recreational and drinking water sources in central North Carolina to assess whether PFAS are present in these fillets that would not be detected by conventional targeted methods. We used liquid chromatography, ion mobility spectrometry, and mass spectrometry (LC-IMS-MS) to collect full scan feature data, performed suspect screening using an in-house library of 100 PFAS for high confidence feature identification, searched for additional PFAS features using non-targeted data analyses, and quantified perfluorooctane sulfonic acid (PFOS) in the fillet samples. A total of 36 PFAS were detected in the fish fillets, including 19 that would not be detected using common targeted methods, with a minimum of 6 and a maximum of 22 in individual fish. Median fillet PFOS levels were concerningly high at 11.6 to 42.3 ppb, and no significant correlation between PFOS levels and number of PFAS per fish was observed. Future PFAS monitoring in this region should target more of these 36 PFAS, and other regions not considered heavily PFAS contaminated should consider incorporating non-targeted analyses into ongoing fish monitoring studies.

**SYNOPSIS:** Concerningly high PFOS levels were detected in fish fillets from recreational and drinking water sources in central North Carolina. Of the 36 total PFAS identified, 19 would not be detected using routine targeted monitoring methods.

**GRAPHICAL ABSTRACT:** 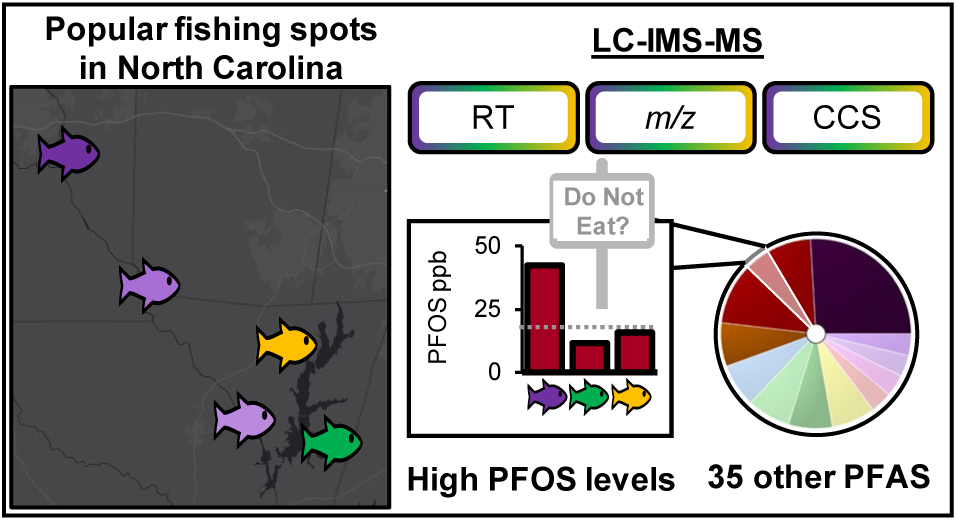

For use in table of contents only

## INTRODUCTION

Per- and polyfluoroalkyl substances (PFAS) are a class of fluorinated anthropogenic chemicals that have been detected in water, soil, and biological tissue worldwide, and PFAS exposure is associated with a suite of adverse health effects. Several recent reviews have detailed how PFAS are manufactured and used,^3, 4^ PFAS global spread and exposure,^5, 6^ and bioaccumulation and toxicity^7, 8^ People are primarily exposed to PFAS through drinking water and food, especially fish consumption.^1, 9^ U.S. federal level regulations on safe PFAS levels in drinking water were enacted for the first time in April 2024.^10^ No federal limits for fish are in place, though several states have implemented their own standards.^1, 11^ Fish consumption advisories are set by comparing measured PFAS concentrations to a reference dose (a daily amount expected to have no adverse effects) to determine how much fish is safe to eat.^2, 12–14^ Establishing reference doses requires toxicity testing to determine what amount of a chemical is safe. These studies typically use validated liquid chromatography separations coupled with targeted mass spectrometry (LC-MS) methods to quantify between 12 and 25 different PFAS, typically including perfluoroalkyl carboxylic acids (PFCAs), perfluorosulfonic acids (PFSAs) including PFOS, and occasionally other types of PFAS.^15^ However, with over 14,000 known PFAS structures and counting,^16^ alternative analytical techniques are essential to understand the full extent of PFAS present in a sample.^15, 17–20^

To increase PFAS coverage in fish tissue analyses, several techniques beyond targeted LC-MS have been used. One option is the total oxidizable precursor assay, which is used in conjunction with targeted MS to assess the presence of precursor PFAS such as fluorotelomers (FTs) and perfluoroalkyl sulfonamides (PFASAs). Pickard et al. quantified two precursor PFAS in fish fillets using this technique, and noted challenges in quantification when precursors were present at low levels.^21^ Another option is non-targeted analyses (NTA) using high-resolution mass spectrometry (HRMS). Rather than pre-selecting specific PFAS to target, NTA techniques rely on accurate mass measurements and other characteristics of PFAS such as fragments, CF_2_ homologous series, and mass defect to filter features and annotate molecules.^15, 22, 23^ NTA may compare feature lists to in-house or publicly-available databases, or propose novel molecule structures when no suitable database match is found. For example, Ren et al. used suspect screening and NTA to detect 15 known and an additional 400 unknown PFAS in freshwater organisms.^24^ Confident annotations of unknowns is a key challenge with any NTA, as Level 1 confidence annotations require matching to a standard.^25, 26^ Ren et al. noted a challenge with low intensity ions that often had no observable MS/MS fragment ions, and thus the unknown PFAS were identified to a Level 4 confidence. Ion mobility spectrometry (IMS) has been suggested as a promising technology to increase annotation confidence^26, 27^ and several studies have demonstrated its utility for PFAS NTA of environmental samples.^28–31^ Another drawback to NTA using HRMS is that these analyses are often qualitative, as it is not possible to create calibration curves for molecules with no existing standards. This can be mitigated by performing a targeted analysis on the sample to quantitate a limited number of PFAS, either on a separate instrument (such as a low resolution mass spectrometer) or concurrently with the high resolution data collection.^15, 32^ However, sample preparation methods that have been validated for use in targeted analyses may not always be optimal for NTA, which favor minimal sample handling to minimize loss of low-abundance analytes. Further, if sample quantities are limited, it may not always be possible to perform both types of analysis.

Fish NTA studies have largely focused on identifying novel PFAS in fish from waters that are highly PFAS contaminated. Little non-targeted effort has been applied in areas considered to have low PFAS levels, despite evidence that water PFAS levels may not be indicative of fish PFAS accumulation.^21^ Routine monitoring of PFAS in fish tissue is not yet commonplace in the U.S., though some states have adopted monitoring programs which test fish tissue using targeted methods to inform consumption advisories; however, the advisories generally only include PFOS.^1^ Therefore, a knowledge gap exists in evaluating whether PFAS that are not commonly targeted are present in fish tissue. Here, we assessed PFAS in fillets of commonly caught and consumed panfish collected from five locations in North Carolina during the summer of 2020 (**Figure 1**). Three sites were in the Haw River, a historically polluted water body under scrutiny for high levels of PFAS contamination.^33^ Two sites were in Jordan Lake, a popular recreational state park and the drinking water source for over 700,000 nearby residents. Water samples simultaneously collected from these locations contained detectable levels of 13 different PFAS, with average PFCA and PFSA concentrations as high or higher than those measured in the Cape Fear River.^34^ However, as of April 2024, no fish consumption advisories for PFAS exist in either the Haw River (HR) or Jordan Lake (JL).^2^ In this study, we analyzed fish fillets on an LC-IMS-HRMS platform and applied target and NTA workflows with the following aims: 1) identify as many PFAS as possible at a high level of confidence; 2) evaluate whether spatial differences in PFAS fingerprints exist between the study sites; and 3) report PFOS concentrations in the fillets to determine whether PFOS levels are a reliable indicator of overall PFAS levels in the fish. Our intent with this non-targeted study was to explore the utility of NTA in fish samples from a lake that is not considered heavily PFAS-impacted, and to inform regulatory testers which PFAS should be targeted in subsequent monitoring efforts in this region.

**Figure 1.**
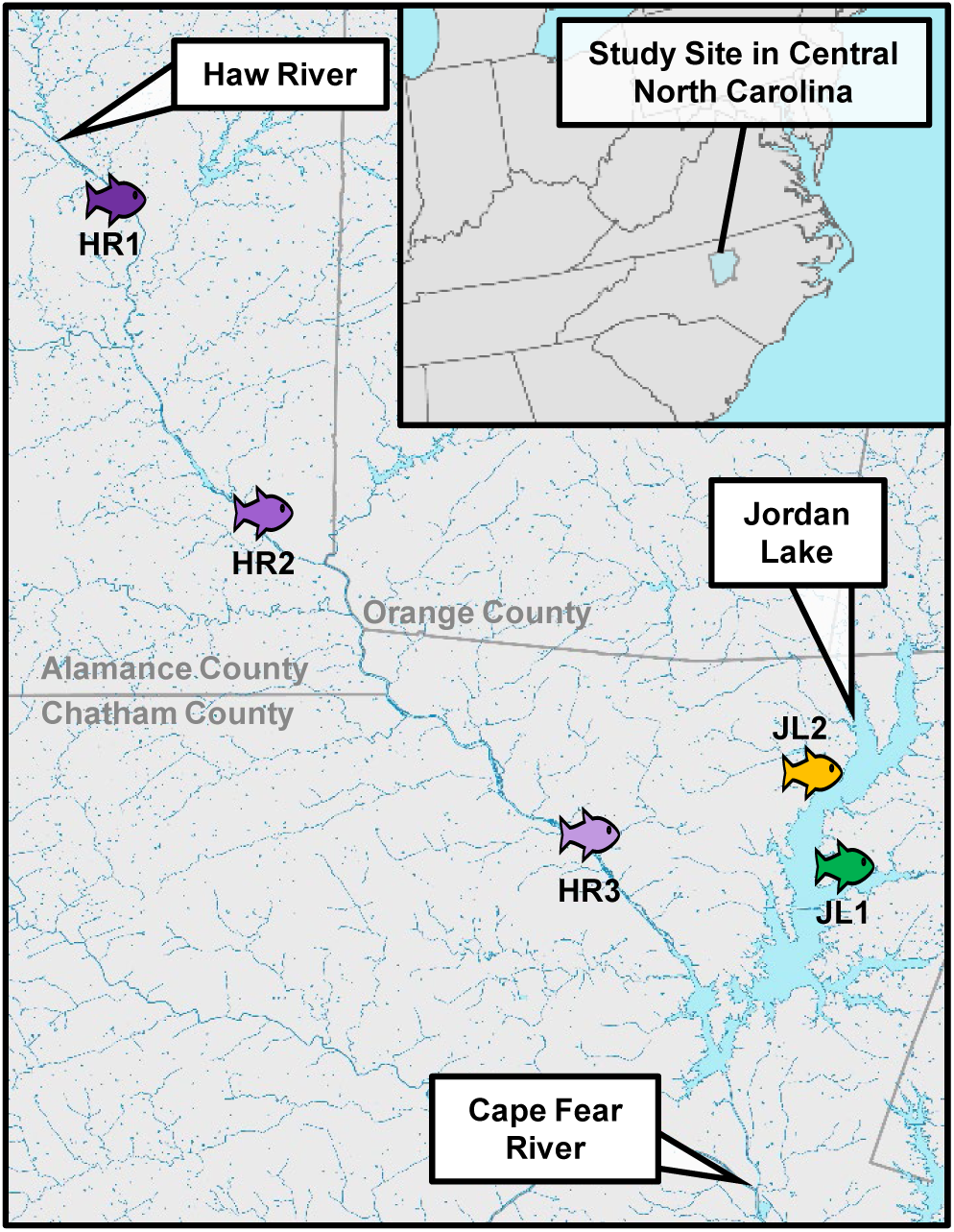
Map of study site showing fish sampling locations. Fish icons show the approximate location of the study site, which spanned three NC counties (Alamance, Orange, and Chatham). Fish were sampled from three sites along the Haw River and two sites within Jordan Lake between May and August 2020.

## MATERIALS AND METHODS

### Study Area and Sample Collection

All animal procedures were performed with approval from the North Carolina State University Institutional Animal Care and Use Committee, protocol number 19-816-01. Fish were collected between May and August 2020 in North Carolina at three sites along the Haw River and two sites in Jordan Lake (**Table 1**, **Figure 1**).

**Table 1.** Sampling sites and number of fish from each site.

| Site code | Site name | n (*Lepomis* spp.) | n (other species) |
| --- | --- | --- | --- |
| HR1 | East Burlington | 13 | 0 |
| HR2 | Saxapahaw | 1 | 1 |
| HR3 | Haw Below 64 | 1 | 0 |
| JL1 | Bell's Landing | 11 | 0 |
| JL2 | Farrington Point | 12 | 9 |

Sampling locations were selected based on proximity to sites with water PFAS data available and angling access. Fish were caught using hook-and-line angling and total length, weight, and sex were recorded. Fillets were removed following accepted decontamination procedures for fish tissue PFAS sampling^41^ and approximately 0.1 to 1 g of skin-on fillet from each fish was stored at -20 °C prior to extraction and analysis. The targeted water analyses for 13 PFAS have been previously reported.^34^ Fish species sampled are detailed in **Table ST1**, and included 38 *Lepomis* spp. sunfish, two white perch (*Morone americana*), four yellow perch (*Perca flavescens*), and four channel catfish (*Ictalurus punctatus*).

### Standards and Reagents

A stable isotope-labeled PFAS standard mix (MPFAC-HIF-ES) was obtained from Wellington Laboratories (Guelph, Canada) containing heavy-labeled PFAS at varying concentrations from 250 to 5,000 ng/ml in methanol. The mix was diluted 1:1 in methanol for use as internal standard. An unlabeled standard mix (PFAC-MXH) from Wellington and farm-raised tilapia fillet purchased from a local grocery store was used for a matrix-matched calibration curve. Optima LC-MS grade methanol, water, and ammonium acetate were used for extractions and mobile phases (Fisher Scientific). The Standard Reference Material 1947 (SRM1947, Lake Michigan Trout) was also obtained from NIST (Gaithersburg, MD) and used for quality control.

### Sample Preparation

In brief, fish fillet samples (0.1 to 1 g wet weight), quality control samples (0.5 g SRM1947), and method blanks (0.5 mL water) were homogenized, spiked with internal standards, and extracted in methanol (5 mL). Clean up of the supernatant included filtration through 0.2 µm nylon syringe filters, dispersive carbon treatment, and centrifugation. The extract was concentrated to dryness in the speedvac and reconstituted in 200 µL of 40% methanol, 60% water by volume buffered with 3 mM ammonium acetate.^42^ Recoveries for 24 PFAS were calculated (**Table ST2**) and were overall quite good, with averages by structural class of 71% for PFCAs, 90% for PFSAs, 91% for FTs, and 70% for PFASAs. Poor recovery was noted for HFPO-DA (24%), N-MeFOSA and N-EtFOSA (4%), and N-MeFOSE and N-EtFOSE (0%); difficult recovery of these 5 analytes have been noted elsewhere.^43, 44^ Detailed sample preparation steps and method characterization including recovery calculations are included in the **Supplemental Methods**.

### LC-IMS-MS Data Collection

Non-targeted sample analysis was performed on an Agilent 6560 IM-QTOF coupled with an Agilent 1290 Infinity UPLC system with consumables and parameters previously described^28^ and detailed in **Tables S1-S4**. In brief, samples were injected in randomized order with a double blank and a QC sample injected between every 10 samples to monitor column carryover and instrument performance. The LC separations were carried out on a C18 column in line with a guard column. Mobile phases included: A) water with 5 mM ammonium acetate, and B) 95% methanol:5% water with 5 mM ammonium acetate. Samples were injected at a volume of 2 µl and a gradient was applied at a flow rate of 0.4 mL per min. An Agilent JetStream electrospray ionization source operated in negative ionization mode was utilized to ionize the samples. Instrument mass calibration was performed by directly injecting Agilent ESI tune mix solution, and tune mix ions were used to convert measured drift times to collision cross section (CCS) values using a single-field calibration.^28^ Ion packets were pulsed through the drift tube using 4-bit multiplexing^47^ and full-scan data was collected in Agilent .d files. Data files were demultiplexed using the PNNL Preprocessor (v 4.0)^48^ with the signal intensity threshold set to 20 counts and 100% pulse coverage and then Agilent IM-MS Browser (v 10.0) was used for single-field CCS calibration.

### Target Screening and Relative Abundance Comparisons

Skyline-daily (v 22.2.1.501) was used for target screening. The target screening workflow uses drift time and retention time filtering against our in-house library of 100 PFAS analytes previously compiled with reference standards.^29^ Identifications required LC retention time alignment with internal standard (as applicable), < 10 ppm *m/z* error versus the library, and CCS filtering with a resolving power window of 30, as well as manual validation of PFAS annotations. Analytes with an exactly matching internal standard were identified at a Level 1a confidence, and all other analytes present in our library were assigned Level 1b.^26, 27^

Following PFAS identification in Skyline, the peak areas of detected PFAS analytes were exported into Excel for further analysis. For analytes detected in method blanks (PFHpA, PFOA, PFOS, and 6:2 diPAP), reporting limits were defined as the average plus 3 standard deviations of the peak area in the method blank, and samples with peak areas below that level were considered not detected. To account for analyte loss from sample extraction and ion suppression in the ESI source, peak areas of the light PFAS analytes were normalized by dividing the peak area of the heavy internal standard. Exact match internal standards were used where possible, and surrogate standards were assigned as needed. Surrogate standards were selected looking first at structural similarity and then at nearest retention time; surrogate assignments are shown in **Table ST3**. Finally, the light-to-heavy peak area ratio was divided by the wet weight of each fillet sample to obtain the relative abundance of each PFAS detected in the fillets (**Equation 1**).

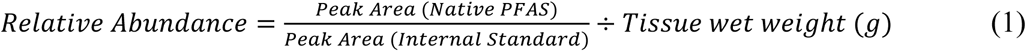

### PFOS Quantitation

PFOS concentrations were calculated using a matrix-matched nine-point calibration curve to correlate light-to-heavy peak area ratios to PFOS concentrations in the sample extracts. The LOD (defined as the average of the blanks plus 3 standard deviations) was 0.0352 ng/ml in the sample extracts (0.388 ng/g in tissue). Three replicates of NIST SRM 1947 were extracted and injected in triplicate to verify calibration curve accuracy against published reference values for PFOS. The average accuracy was 101% with a 10.2% standard deviation. Samples were originally run without the calibration curve, and then later re-run with the calibration curve; unfortunately, due to concerns with sample degradation during storage, only PFOS could be confidently quantified. Detailed methods on the quantification and verification of data reproducibility are included in the **Supplemental Methods, Figure S1,** and **Table ST4**. Although additional analytes could not be reliably quantified in this study, PFOS is the most relevant PFAS to human exposure via fish consumption given that state consumption advisories exist are typically limited to PFOS only, and we believe reporting these results provides important context for fish PFOS concentrations that have not been previously evaluated. Future NTA studies relevant to human PFAS exposure should include absolute quantification of the most relevant PFAS to continue increasing our understanding of human PFAS exposure routes.

### Non-Targeted Data Analysis

Agilent MassProfiler (v 10.0) was used to generate a feature list for all fish samples, filtering the demultiplexed data files by ion intensity (≥ 100 counts), RT tolerance (± 0.3 min), DT tolerance (2.0%), mass tolerance (± 10.0 ppm + 2.0 mDa), and abundance compared to method blanks (fold change ≥ 2.0). Fluoromatch-IM (v. 1.2) was then used to search the feature list for additional PFAS not present in our library. Fluoromatch-IM flags CF_2_(n) homologous series and allows the user to filter the data based on other PFAS characteristics such as mass defect, negative Kendrick mass defect (KMD), and drift time and retention time ordering. Features determined to be possible PFAS were added to the Skyline document for visualization and manual verification.^49^ Additionally, using results from the initial target screening, molecular formulas for PFAS not present in our library but theoretically part of a CF2(n) homologous series with detected analytes were added to the Skyline document. The CCS was measured for features detected in at least one sample and annotations were assigned using the criteria described below, adapted from previously published guidance on confidence reporting for PFAS (which does not include the IMS dimension)^26^ and for ion mobility data (which does not include PFAS characteristics like homologous series and mass defect).^27^ All identifications Level 3 and higher required < 10 ppm *m/z* error and retention time ordering consistent with in-house library values. Level 1a and 1b required matching to a reference standard as described in the target screening methods. Level 2a required a CCS < 2% in comparison to external database values. Level 2b required a fit to in-house CCS vs *m/z* trendlines and presence of 1 or more analyte in homologous series identified at Level 1 confidence. Level 3a and 3b required the same criteria as 2a and 2b, respectively, but contained a structural element that could not be elucidated (e.g. branching position). Analytes annotated using these non-targeted methods were added to the Skyline document for peak picking and relative abundance comparisons as described above. We planned to assign level 5 to features annotated using Fluoromatch-IM that did not match any database CCS and *m/z* entry; however, none were ultimately found so this level was not used.

### Statistical Analysis

Statistical analyses were conducted in R (v.4.2.1) and GraphPad Prism (v.10.2.2). Prior to analysis, PFAS relative abundances were normalized by first adding a pseudo count to ensure there were no zeros, and then applying a log_2_ transformation. Principal component analysis (PCA) was performed using the function ‘prcomp’ and results were visualized using ggplot2. Differences between sampling timepoints were assessed using multivariate analysis of variance (MANOVA) using the function ‘manova’. Differences in sampling locations were visualized using a heatmap, which was constructed using the function ‘pheatmap’. Hierarchical clustering was performed on samples and analytes using the Euclidean distance metric and average linkage.^50^ The Toxicological Prioritization Index (ToxPi) tool (v 2.3)^51^ was used for visualization of relative abundance of PFAS from individual fish or average abundance from different sample groups. Relationships between continuous variables (fish length, number of PFAS detected) and PFOS concentration were assessed using regression analysis, and significance of the slope was evaluated using the t-test for regression coefficients. Relationships between sex, site, and PFOS concentration were assessed using a two-way ANOVA with Šídák’s multiple comparisons test. Relationships between site and PFOS concentration were assessed using one-way ANOVA with Tukey’s multiple comparisons correction.

## RESULTS AND DISCUSSION

### High Confidence PFAS Identifications

All 48 fillets were analyzed using LC-IMS-MS to collect full scan feature data which was screened against our library of 100 PFAS, and additional PFAS were annotated based on mass defect, presence in homologous series, LC retention time ordering, IMS drift time ordering, and *m/z* vs CCS trend lines (**Figure S2**). In total, 36 PFAS analytes were identified at high confidence in at least one fish fillet, including 27 PFAS at a Level 1a or 1b confidence, 2 PFAS at Level 2b, 6 PFAS at Level 3a, and 1 PFAS at Level 3b (**Tables 2, ST5, ST6**). Because the exact position of branching for the PFSA-br isomers^3^ could not be elucidated^28^, the following discussion counts linear and branched PFSA isomers as one analyte. The maximum number of PFAS analytes detected in one fish was 22, while the minimum was 6. Notably, 5 analytes (PFNA, PFUdA, PFDA, PFOS, and PFDS) were detected in every fish. Using Mass Profiler, 15,783 LC-IMS-MS features were found. Fluoromatch-IM’s algorithm was applied to the feature list (**Table ST7**) and features not matching our filtering criteria for possible PFAS (negative mass defect, negative Kendrick mass defect, retention time ordering, drift time ordering, presence in homologous series) were removed. After removing features previously identified in our target screening workflow, 55 features remained, but none were annotated as PFAS. Investigation of these peaks in Skyline revealed that 53 features were smearing or noise (**Figure S3**) while the remaining 2 features were incorrectly ordered with others in the flagged homologous series and did not match any formula listed in the CompTox Chemical Dashboard. While it is possible that deeper investigation could uncover additional and potentially novel PFAS, our primary aim was to investigate what PFAS we could confidently identify in these fillets. Any additional annotations using this workflow could not be higher than Level 5 confidence. As NTA tools improve, this data set can be revisited and mined to determine if additional PFAS can be discovered.

**Table 2.** All PFAS identified in at least one *Lepomis* fish fillet. PFAS are grouped by structural class. Confidence levels are based on the Schymanski scale modified by Charbonnet *et al.* for PFAS use.

| Structural class | Acronym | Full name | Confidence | Total detection frequency |
| --- | --- | --- | --- | --- |
| PFCA | PFOA<sup>a, b</sup> | perfluorooctanoic acid | 1a | 8% |
|  | PFNA<sup>a, b</sup> | perfluorononanoic acid | 1a | 31% |
|  | PFDA<sup>a, b</sup> | perfluorodecanoic acid | 1a | 100% |
|  | PFUdA<sup>b</sup> | perfluoroundecanoic acid | 1a | 100% |
|  | PFDoA<sup>b</sup> | perfluorododecanoic acid | 1a | 100% |
|  | PFTrDA<sup>b</sup> | perfluorotridecanoic acid | 1b | 94% |
|  | PFTeDA<sup>b</sup> | perfluorotetradecanoic acid | 1a | 88% |
|  | PFPeDA | perfluoropentadecanoic acid | 2b | 46% |
|  | PFHxDA | perfluorohexadecanoic acid | 1b | 13% |
| PFSA | PFHxS-br | perfluorohexanesulfonate (branched) | 3a | 4% |
|  | PFHxS<sup>a, b</sup> | perfluorohexanesulfonate | 1a | 73% |
|  | PFHpS<sup>b</sup> | perfluoroheptanesulfonate | 1b | 42% |
|  | PFOS-br | perfluorooctanesulfonate (branched) | 3a | 100% |
|  | PFOS<sup>a, b</sup> | perfluorooctanesulfonate | 1a | 100% |
|  | PFNS-br | perfluorononanesulfonate (branched) | 3a | 31% |
|  | PFNS<sup>b</sup> | perfluorononanesulfonate | 1b | 58% |
|  | PFDS-br | perfluorodecanesulfonate (branched) | 3a | 100% |
|  | PFDS<sup>b</sup> | perfluorodecanesulfonate | 1b | 100% |
|  | PFUdS-br | perfluoroundecanesulfonate (branched) | 3b | 6% |
|  | PFUdS | perfluoroundecanesulfonate | 2b | 10% |
|  | PFDoS-br | perfluorododecanesulfonate (branched) | 3a | 2% |
|  | PFDoS<sup>b</sup> | perfluorododecanesulfonate | 1b | 31% |
|  | Cl-PFOS | chloro-perfluorooctanesulfonic acid | 1b | 10% |
| FT | 8:2 FTS<sup>b</sup> | 8:2 fluorotelomer sulfonic acid | 1a | 2% |
|  | 8:2 FTPA | (1H,1H,2H,2H-Heptafluorodecyl)phosphonic acid | 1b | 2% |
|  | 10:2 FTS | 10:2 fluorotelomer sulfonic acid | 1b | 2% |
| PFASA | FBSA | perfluoro-1-butanedisulfonamide | 1b | 44% |
|  | FHxSA | perfluoro-1-hexanedisulfonamide | 1b | 4% |
|  | FOSA<sup>b</sup> | perfluoro-1-octanedisulfonamide | 1a | 48% |
|  | N-MeFOSAA<sup>b</sup> | N-methylperfluoro-1-octanedisulfonamidoacetic acid | 1a | 13% |
|  | N-EtFOSAA<sup>b</sup> | N-ethylperfluoro-1-octanedisulfonamidoacetic acid | 1a | 2% |
| PFP | 6:6 PFPi | 6:6 Perfluorophosphinic acid | 1b | 27% |
|  | 6:8 PFPi | 6:8 Perfluorophosphinic acid | 1b | 35% |
|  | 6:2 diPAP | bis(1H,1H,2H,2H-perfluorooctyl)phosphate | 1b | 10% |
|  | 6:2/8:2 diPAP | 6:2/8:2 Fluorotelomer phosphate diester | 1b | 4% |
|  | 8:2 diPAP | bis(1H,1H,2H,2H-perfluorodecyl)phosphate | 1b | 2% |

<sup>a</sup>: previously detected in water
<sup>b</sup>: targeted in quantitative method EPA 1633
-br: branched isomer

PFAS have previously been reported present in Haw River and Jordan Lake water, and a recent study targeted and detected 13 different PFAS in water samples taken from the exact same sampling locations and time points as the fish samples.^34^ Of the 13 PFAS previously reported in water samples, 5 (PFOA, PFNA, PFDA, PFHxS, and PFOS) were also detected in our fish samples. These 5 analytes are all long-chain PFAS^3, 52^ which are known to bioaccumulate in tissue.^7, 18, 53^ However, 8 other PFAS (PFBA, PFPeA, PFHxA, PFHpA, PFBS, 4:2 FTS, 6:2 FTS, and HFPO-DA (“GenX”)) were detected in the targeted water study but not in the fillets, despite all 8 of these being present in our in-house library. These analytes were all shorter chain PFAS that are known to have shorter biological half-lives, so it is possible that these analytes had not accumulated to detectable levels in the fillets. A key limitation of this comparison is that water and fish analyses were conducted using separate targeted and non-targeted approaches. The longer chain PFAS most frequently detected in the fish samples were not targeted in the water analysis of the Haw River and Jordan Lake, so it is unknown whether they would have been detectable in the water. Future studies aiming to understand differences in water and fish tissue PFAS presence should analyze both sample types on both platforms. Despite these limitations, our study provides evidence that water PFAS levels do not accurately describe fish PFAS accumulation, which is also supported by other researchers.^18, 21, 54^

### Spatial PFAS Trends

To evaluate differences in PFAS accumulation in fish fillets at the different sampling sites, detection frequencies and relative abundances of the PFAS identified at Level 3 or higher were compared. Of the fish species collected, the majority (36 samples) were *Lepomis spp.* sunfish from sites HR1, JL1, and JL2. Due to the low number of samples from the other sites and species, subsequent data visualizations and statistical analyses include only the PFAS detected in at least one *Lepomis spp.* sample to minimize confounding effects from biological variation. Results from sites HR2 and HR3 and the other fish species are included in the SI. The sunfish were caught over four months in the summer of 2020. A statistical analysis by MANOVA of relative abundances of the 7 PFAS detected in 90% or more of the individual sunfish samples showed no sampling date differences (p-value = 0.249, **Figure S4**). This result was not surprising because these analytes are primarily long chain PFAS known to have long biological half-lives. Therefore, subsequent analysis combined all sunfish from each site regardless of the sampling date.

Unsurprisingly, the HR1 sunfish had more PFAS analytes detected than the JL1 and JL2 sites (**Figure 2**), with 25 PFAS detected at HR1, 15 PFAS at JL1, and 20 PFAS at JL2. While some of these PFAS were detected in only 1 fillet, we have opted to include them in this discussion due to the high confidence that the IMS dimension affords these identifications. The average number of PFAS detected per fish followed the same spatial trend, with 15.4 PFAS per fillet on average at HR1 (range: 10 to 22), 8.4 PFAS per fillet at JL1 (range: 6 to 11), and 10.3 PFAS per fillet at JL2 (range: 7 to 17). Targeted methods for PFAS in fish fillet monitor for a predefined list of *m/z* values, for example, EPA 1633 tests for 40 PFAS. Of the 40 PFAS on the 1633 target list, 17 were detected in the fish fillets. The remaining 13 PFAS identified in the fish would not be detected using common targeted methods. The HR1 site is immediately downstream of a wastewater treatment plant, which is a known point source of PFAS into the river, and so we expected to find more PFAS using NTA at HR1 compared to JL1 or JL2. Nevertheless, PFAS were detected using NTA at both JL1 and JL2. For example, 100% of sunfish fillets from HR1 and 17% from JL2 had detectable levels of PFPeDA. PFPeDA is a PFCA with 15 carbons which is not targeted in any validated regulatory method and has rarely been reported in literature, possibly due to how challenging it is to obtain a standard for this molecule. Additionally, more than 40% of sunfish fillets at all 3 sites had detectable FBSA, a short chain sulfonamide PFAS which is not commonly targeted in fish or water methods despite standards being available.

**Figure 2.**
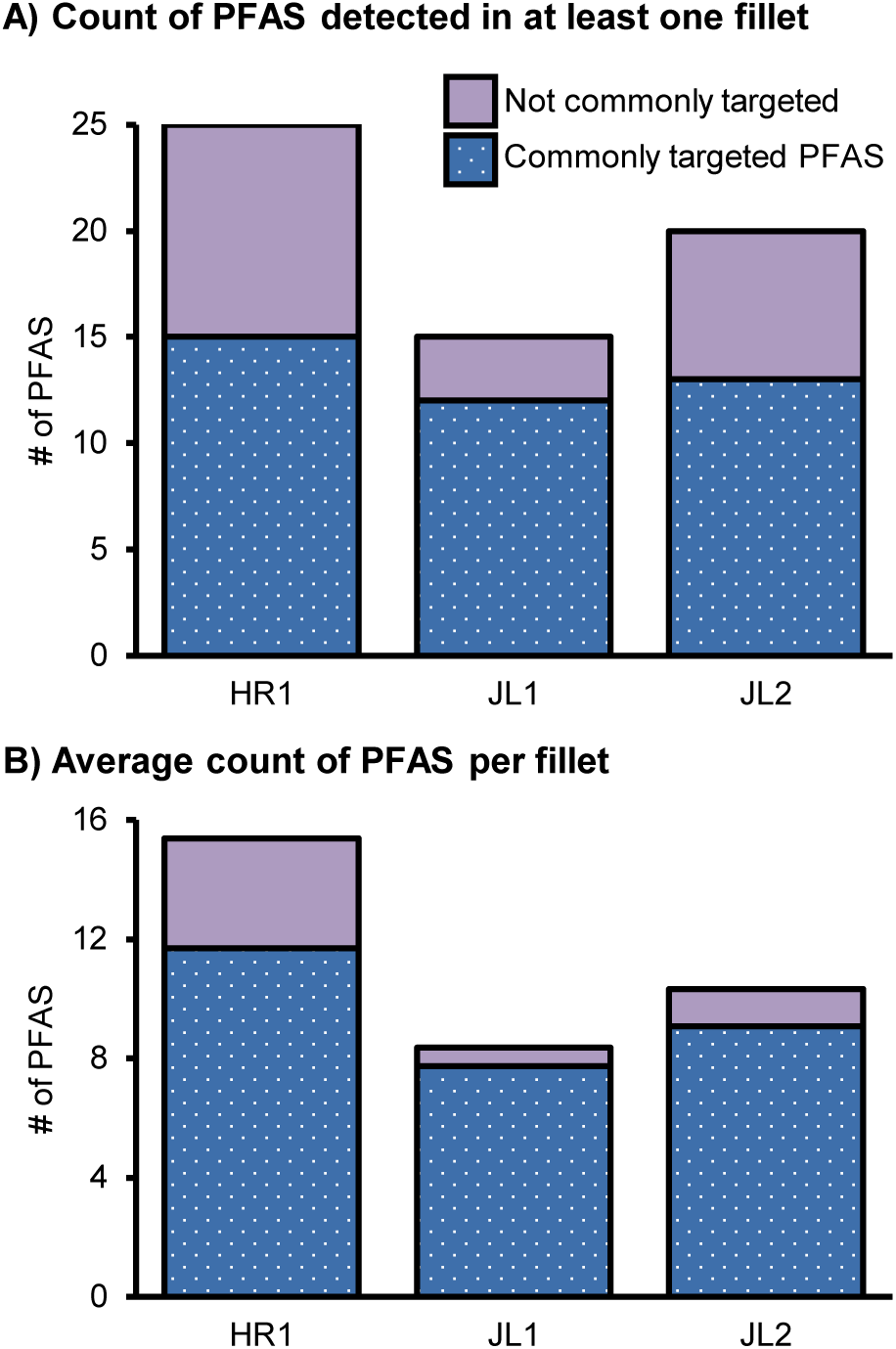
Number of PFAS detected **A)** in at least one *Lepomis* fish fillet and **B)** on average in fillets from the three sampling sites. The dotted dark blue bars are PFAS that are commonly targeted in validated methods such as EPA 1633. The solid light purple bars are PFAS that were uniquely detected using non-targeted analysis.

The relative abundances of PFAS in the sunfish fillets (**Equation 1, Table ST4**) were compared using ToxPi to visualize PFAS analyte abundances at each site.^51^ In ToxPi, each analyte is represented by a slice, and the relative abundance of each PFAS is normalized to the maximum abundance of that PFAS detected in a sample, so a full-length slice represents the maximum measured abundance of that specific analyte in the sample set. The width of each slice is proportional to the number of analytes grouped within the slice; here, they are grouped by structural class and whether the molecule is commonly targeted. In general, sunfish from HR1 had higher relative abundances of all detected PFAS, as expected. The relative abundances of the commonly targeted PFAS were generally much lower in fillets from JL1 and JL2. Principal component analysis and heatmap clustering showed that HR fillets separated from JL fillets due to the higher abundances of the PFAS detected with > 70% frequency (the C10-C14 PFCAs and the C6, C8, and C10 PFSAs) in the HR fish (**Figure S5, S6**), though separation of the two JL sites was not observed. However, some notable observations were made in individual fillets (**Figure 3**). One fillet from JL1 had the highest relative abundance of 6:2 diPAP and 8:2 diPAP of all fish tested, despite having some of the lowest reported PFOS concentrations. Phosphate-containing PFAS are not commonly targeted despite standards being available, and one recent study suggests that their levels are grossly underestimated in most targeted groundwater studies.^20^ PFUdS, the 11-carbon PFSA, is not commonly targeted and to our knowledge has not previously been reported detected in fish. Linear and branched isomers of PFUdS were detected in 23% of sunfish from HR1 as well as in one fish from JL2. This JL2 fish had a relative abundance of PFUdS nearly as high as the highest fish from HR1, despite having comparatively lower abundances of the other PFSAs. Toxicity information is not available for most of these compounds, so it is not clear how much risk to health consumption of these fish could pose.

**Figure 3.**
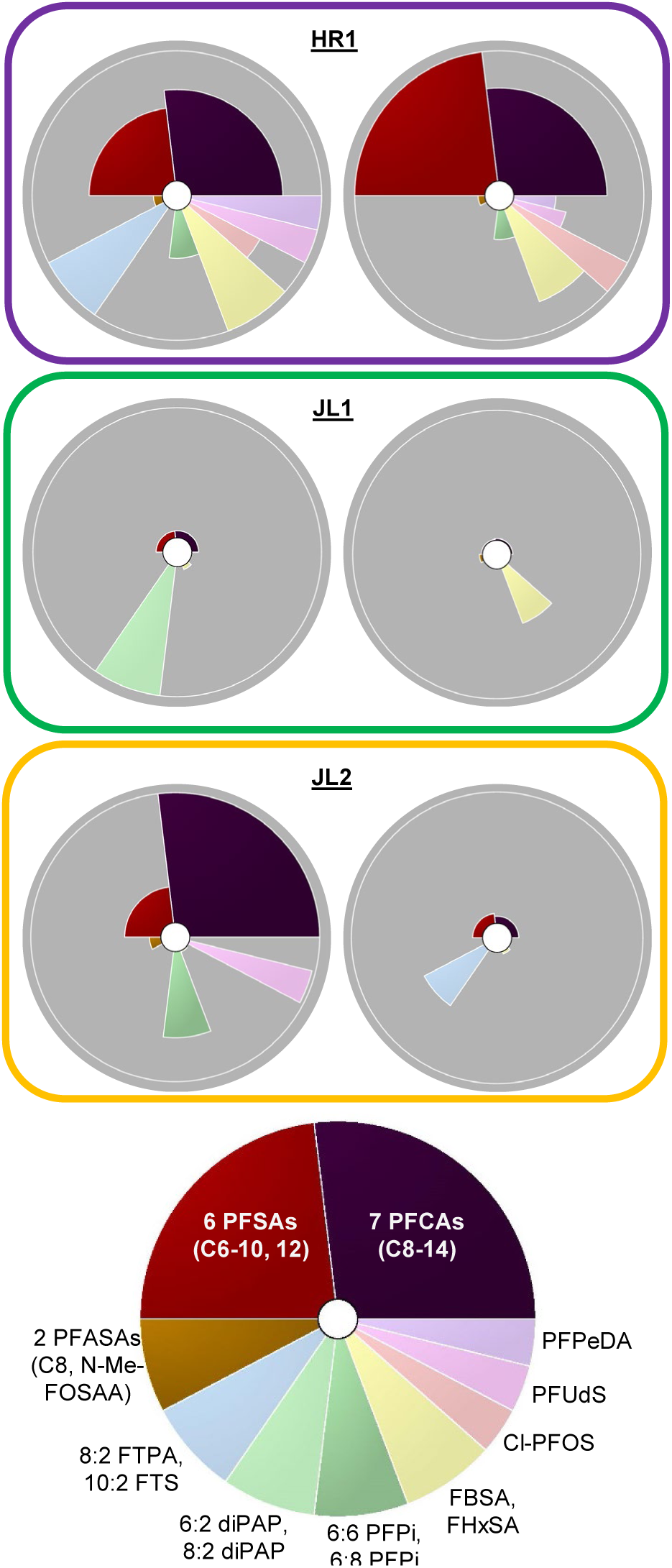
ToxPi visualizations showing relative PFAS abundances in two individual *Lepomis* fillets from each site. The commonly targeted PFAS are shown in dark colors while the PFAS not commonly targeted are shown in pastels. Structurally similar PFAS are combined into one slice and each slice is weighted based on the number of PFAS it represents. The length of each slice indicates the abundance of that PFAS relative to its abundance in all other samples so that a full-length slice represents the maximum abundance of that molecule.

### PFOS Concentrations

PFOS concentrations in the *Lepomis spp.* sunfish fillets ranged from 7.5 to 124 ppb, with the highest at HR1 (median 42.3 ppb), followed by JL2 (15.6 ppb) and JL1 (11.6 ppb) (**Figure 4, Table ST4)**. The differences between the HR samples and the JL samples were statistically significant by one-way ANOVA comparisons using Tukey’s multiple comparisons test (F (2, 33) = 18.57, adjusted p-value < 0.0001). The JL1 and JL2 sites were not significantly different from each other (adjusted p-value = 0.700). Sex and fish length were evaluated as potential covariates for PFOS concentrations (**Figures S7, S8**). No sex differences were observed at JL1 or JL2, and half of the fish from HR1 were not sexed; therefore, fish from each site were grouped regardless of sex. There was a positive correlation between fish length and PFOS concentration at JL1 and JL2, but not at HR1. Because fish length is related to age within a population^62^, we expected to see a positive correlation between length and PFOS levels due to the long biological half-life of PFOS in muscle tissue. However, numerous factors unrelated to PFOS exposure, such as pharmacokinetic changes from other chemicals present at HR1 and differences in the riverine compared to lake ecosystems, could explain this lack of relationship observed in the HR1 fish.^63^ These results suggest that consuming larger sunfish could result in greater PFOS exposure; however, at sites with high water PFOS levels, such as HR1, smaller fish may not necessarily have lower levels of PFOS than larger fish. Additional studies are needed to investigate this observation further.

**Figure 4.**
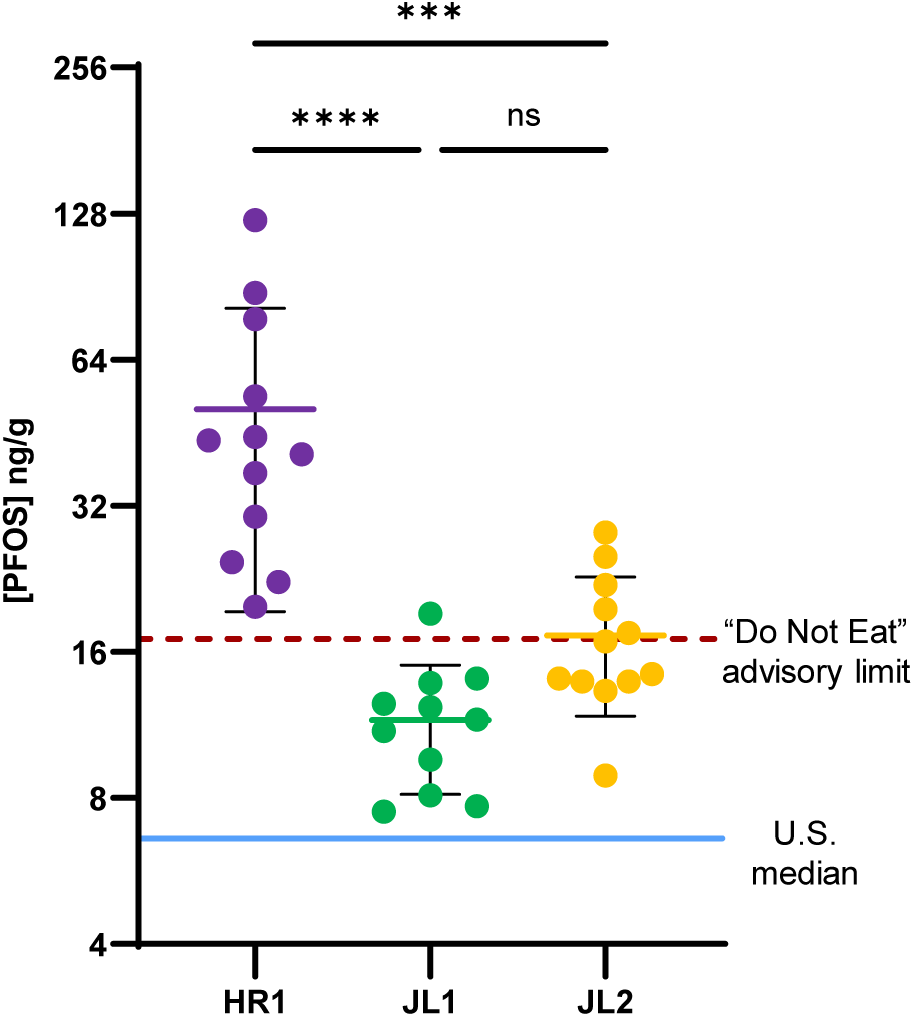
Concentration of PFOS measured in fillets from each site. The blue horizontal line indicates the national median levels for freshwater fish across the U.S.^1^ The dashed red line at 17.2 ppb corresponds to a “Do Not Eat” level of PFOS based on the only fish consumption advisory currently in effect in North Carolina, which applies to the lower Cape Fear River only.^2^

To our knowledge, these are the first PFOS concentrations reported in fish from Jordan Lake and the upper Haw River. A recent EPA study of 44 species of freshwater fish sampled between 2013 and 2015^1^ reported that the median PFOS level in fish from rivers and streams across the U.S. was 6.60 ppb. The median PFOS concentrations we report here in the *Lepomis spp.* sunfish fillets are higher than reported by the EPA study in all cases. In the geographically closest study we could find, a 2007 study of North Carolina fish, Delinsky et al. reported a median concentration of 30.3 ppb PFOS (range: 15.9 to 47.5 ppb) in bluegill sunfish collected from the Haw River, south of the inflow to Jordan Lake.^64^ The median PFOS concentrations we report in fish from the upper Haw River are higher than those reported downstream in 2007, while the Jordan Lake fish had generally lower median PFOS concentrations that were nevertheless within the observed range from this area in 2007. More recently, the North Carolina Department of Health and Human Services reported average PFOS concentrations of 23.0 ppb in bluegill from the middle and lower Cape Fear River in the summer of 2022, which is high enough for a “do not eat” advisory which is triggered at levels exceeding 17.2 ppb. The PFOS concentrations we measured in Jordan Lake fillets were comparable to the advisory triggering levels in the Cape Fear, and levels in the Haw River fillets were more than twice that high. While the Cape Fear River has garnered a lot of attention due to GenX contamination from Chemours and the Haw River is the subject of ongoing litigation around PFAS pollution, Jordan Lake is not considered a highly contaminated area and has not been as highly scrutinized. Our results provide evidence that fish in this watershed have concerningly high levels of PFOS, and urgent attention is needed in this area.

Currently in North Carolina, PFOS is the only PFAS applicable for fish consumption advisories. However, we found a total of 36 different PFAS, including PFOS, in fish fillets from Jordan Lake and the Haw River. To evaluate whether PFOS concentrations in fish fillets are associated with greater numbers of different PFAS detected, regression analysis was performed between PFOS concentrations and number of PFAS detected in sunfish fillets from each site. Surprisingly, PFOS concentration did not generally appear to correlate with increasing number of PFAS detected (**Figure 5**). The goodness of fit as measured by R^2^ was low at all three sites, and a statistically significant correlation was observed at JL2 (p = 0.03) but not at HR1 (p = 0.50) or JL1 (p = 0.62). These results indicate that there is not a consistent relationship between PFOS concentration and number of different PFAS in sunfish fillets. Notably, the sunfish with the lowest PFOS concentration at JL1 (7.50 ppb) had the highest number of PFAS detected at that site (11). This fish had some of the highest FBSA and FHxSA levels of all the sunfish fillets. At HR1, the sunfish with the lowest PFOS concentration (19.8 ppb) had the third-highest number of PFAS detected (19), including the long chain PFCAs PFPeDA and PFHxDA. All the long-chain PFCAs and PFSAs are thought to be harmful and some of them are more bioaccumulative than PFOS. In North Carolina as well as in most other states in the U.S., however, water and fish are rarely tested for these long-chain PFAS. Our results suggest that they could be present at harmful levels in fish even when PFOS levels are considered safe. It is possible that the same trends would not be observed at lower PFOS concentrations, which we could not evaluate here because our sample set did not include any fish with low enough PFOS levels to be safely consumed more than one time per year based on the reference dose applied in the Cape Fear River.

**Figure 5.**
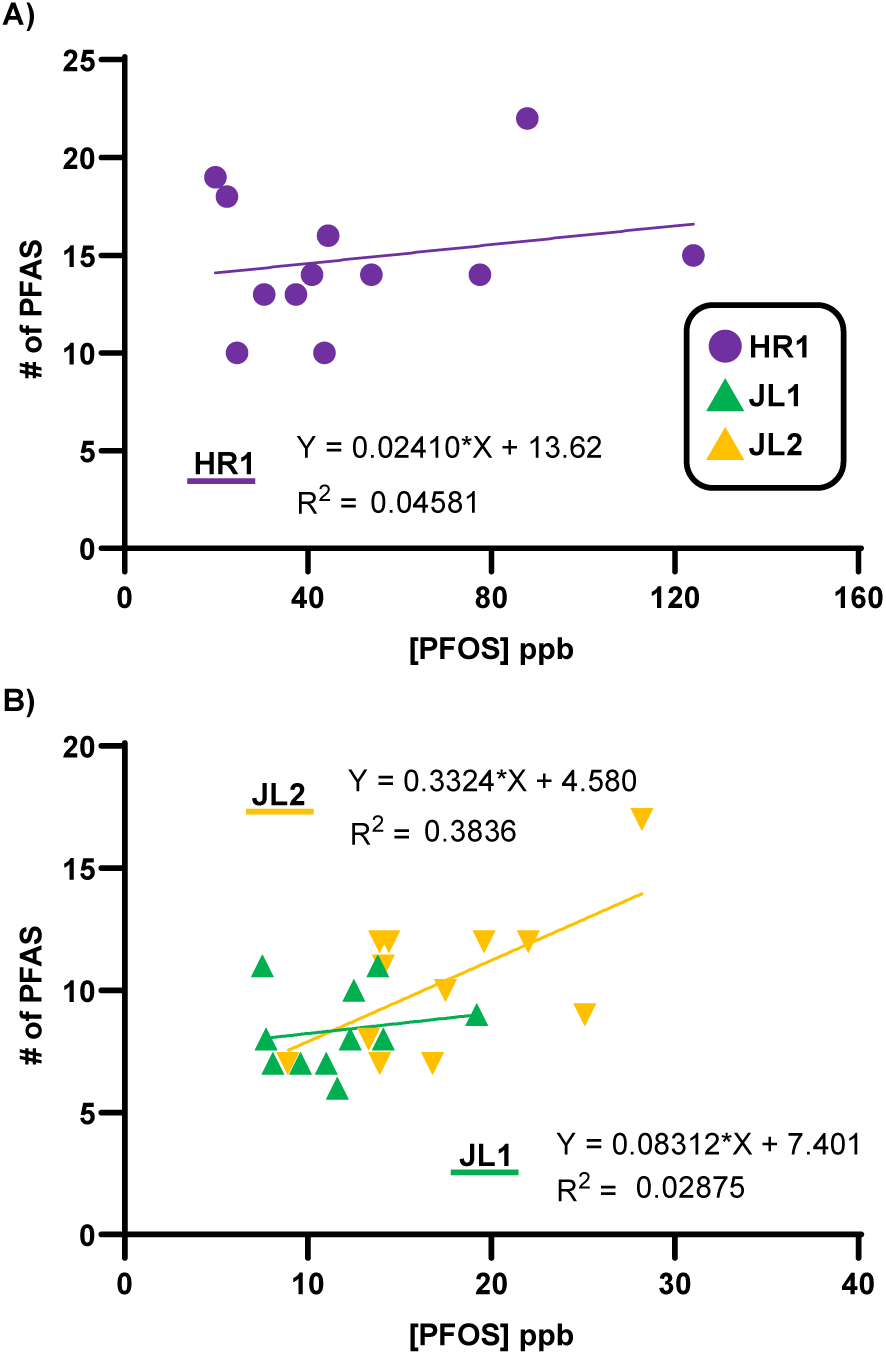
Concentration of PFOS measured versus number of PFAS detected in individual *Lepomis* fillets from **A)** the Haw River site and **B)** the Jordan Lake sites. A general trend of increasing [PFOS] with number of PFAS detected was observed. Regression analysis indicated a statistically significant association at JL2 (p = 0.03) but not at JL1 or HR1.

### Implications

In this study, NTA was performed to find PFAS in fish from the Haw River and Jordan Lake in North Carolina, a previously unstudied water body upstream of the Cape Fear River (which has received considerable attention for its PFAS contamination). In total, 36 different PFAS were identified in fish fillets at a high level of confidence made possible by multidimensional LC-IMS-MS separations. Of these 36 PFAS, 5 have been previously reported in the water from these sites. Only 17 of these 36 PFAS are targeted in the validated EPA method for PFAS in biological samples, EPA 1633. The remaining 19 PFAS are not commonly targeted in fish or water studies, and therefore little information is available on their toxicities and environmental presence. While these 19 PFAS were not quantified in this study, standards are available for most of them and future fish monitoring studies in this region should include calibration curves for as many of these PFAS as feasible. Our unknown discovery workflow flagged 55 additional features as possible PFAS; however, manual inspection of the peaks determined that they were not likely PFAS and could not be annotated. Incorporating more CCS values into existing databases will help increase the number of PFAS that may be confidently identified and subsequently tracked and monitored in future studies.

PFOS concentrations in all fillets exceeded the “no more than one meal per year” consumption advisory limit which is triggered at 1.4 ppb in the lower Cape Fear River, with the lowest at 7.50 ppb and the highest at 124 ppb. To our knowledge, no PFAS fish monitoring is in place in North Carolina outside of the Cape Fear River. The fish in our study were collected in 2020, so it is possible that PFAS levels today are different from those we report here. Given that conventional wastewater treatment methods do not remove PFAS and that very little federal or state-level PFAS regulations are in place, we urge government and regulatory agencies to implement a fish PFAS monitoring program for this region so the millions of annual visitors to Jordan Lake State Park are informed of any potential risk of PFAS exposure they might accept from consuming fish from this lake. While our study area was limited to a small geographic region in North Carolina, we believe our results provide clear evidence that fish PFAS monitoring studies should include NTA even in areas not considered heavily PFAS-impacted.

## ASSOCIATED CONTENT

### Supporting Information

Fish fillet sample preparation methods; Consumables used in LC-IMS-MS analysis (Table S1); LC gradient settings (Table S2); Electrospray ionization source settings (Table S3); IMS-MS settings (Table S4); Absolute and semi-quantification methods; Reproducibility of PFOS calculations (Figure S1); Accuracy and precision of PFAS quantitation (Table S5); CCS vs *m/z* trendline example (Figure S2); Example of possible PFAS ruled out (Figure S3); PCA of sampling date (Figure S4); PCA of sampling location (Figure S5); Heat map by location (Figure S6); Sex differences in PFOS concentration (Figure S7); Fish length and PFOS concentration (Figure S8) (PDF)

### Supplemental Tables

Fish sample information (ST1); Efficiency and recovery (ST2); PFAS identifications (ST3); PFOS quantitation (ST4); Normalized areas (ST5); QC normalized areas (ST6); FM-IM results (ST7) (XLSX)

### Panorama

Skyline files with all LC-IMS-MS data are available at: https://panoramaweb.org/2023-NC-Fish-PFAS-NTA.url

## AUTHOR INFORMATION

The authors declare no competing interests.

## ACKNOWLEDGEMENTS

This work was funded by grants from the National Institute of Environmental Health Sciences (P42 ES027704 and P42 ES031009), and a cooperative agreement with the Environmental Protection Agency (STAR RD 84003201). The views expressed in this manuscript do not reflect those of the funding agencies. AKB wishes to thank Philip M. McDaniel of the UNC-Chapel Hill Libraries for his help in generating the map shown in Figure 1; Kaylie Donelson, Guozhi Zhang, Jack Ryan, Amie Solosky, Ashlee Falls, and Greg Kudzin for their valuable feedback on early versions of the figures and manuscript; and Kara Joseph for assistance with sample preparation.

