## Supporting Information Document for "Assessing Per- and Polyfluoroalkyl Substances (PFAS) in Fish Fillet Using Non-Targeted Analyses"

**Fish fillet sample preparation methods.**

Fish fillet samples, quality control samples (0.5 g SRM1947), and method blanks (0.5 mL water) were homogenized in a bead mill (2 mL polypropylene tubes, Fisher Scientific, part number 15-340-152). Sample wet weights for the fish fillets ranged from 0.1 to 1 g (**Table S1**). The entire sample was extracted using the following procedure and the range of sample weights was accounted for in data analysis. The homogenized tissue was spiked with internal standards (5 µL MPFAC-HIF-ES diluted 1:1 in methanol) and extracted in methanol (5 mL) in polypropylene tubes (15 mL, Falcon, 352196). Samples were sonicated for approximately 1 h and then centrifuged at 4 °C and 10,000 rpm for 5 min. The supernatant was decanted into polypropylene syringes (5 mL, MilliporeSigma, Z683582) and filtered through 0.2 µm nylon syringe filters (25 mm, Whatman, 6751-2502) into new polypropylene tubes loaded with 10 mg dispersive carbon (Envicarb, 57210). Samples were shaken and inverted by hand for 1 min and centrifuged again. The supernatant was transferred into new polypropylene tubes and concentrated to dryness in the speedvac at 45 °C and full vacuum. Samples were reconstituted in 200 µL of 40% methanol, 60% water by volume buffered with 3 mM ammonium acetate, using the methanol-first method described by Weed et al. [1] in which the methanol portion is added to the sample first, vortexed, and then the aqueous portion is added. Reconstituted samples were transferred to polypropylene LC vials (Agilent 1 ml vials, 5182-0567, and 11 mm caps, 5182-0542) with inserts (Agilent 250 µl inserts, 5182-0549).

Method performance was evaluated using both process efficiency and recovery [2, 3]. Fish tissue was spiked with a heavy-labeled standard mix (5 µl MPFAC-HIF-ES diluted 1:1 in methanol) either before or after being extracted as described above. Vials of neat solvent matching the reconstitution buffer (40% methanol, 60% water, with 3 mM ammonium acetate) were spiked with the same standard mix to the same concentration. Recovery was calculated with **Equation S1**, and process efficiency was calculated with **Equation S2**. The recovery percentage indicates how much analyte is recovered during the sample extraction and clean up steps, while process efficiency accounts for both recovery and matrix effects.

$$Recovery = \frac{Peak\ Area\ (pre\text{-}extraction\ spike)}{Peak\ Area\ (post\text{-}extraction\ spike)} \times 100\% \qquad (S1)$$

$$Process\ Efficiency = \frac{Peak\ Area\ (pre\text{-}extraction\ spike)}{Peak\ Area\ (spiked, unextracted\ neat\ solvent)} \times 100\% \qquad (S2)$$

Recoveries ranged from 0% to 97% with an average of 62% for all analytes (**Table ST2**). The analytes with recoveries below 30% included the two shortest-chain PFCAs (PFBA and PFPeA), the perfluoroether carboxylic acid HFPO-DA ("GenX"), two sulfonamides, and two sulfonamido ethanols. Poor recovery of short chain PFCAs and PFECAs and poor method performance for these sulfonamide compounds has been reported elsewhere. All other analytes had recoveries of ~70% or greater, which is considered adequate.

Process efficiencies ranged from 0% to 430% with an average of 36% for all analytes (**Table ST2**). While no specific criteria exist for process efficiency, in comparison with recovery, it provides valuable information about matrix effects. Most analytes had process efficiencies below 30%, which is likely indicative of ionization suppression due to biomolecules present in the fish matrix. PFTeDA had a surprisingly high process efficiency at 430%, which could be due to ionization enhancement or increased solubility when fish matrix is present.

### Table S1. Consumables used in LC-IMS-MS analysis

| Item | Size | Manufacturer | Part number |
|---|---|---|---|
| C18 column (Zorbax Eclipse Plus) | 2.1 x 50 mm, 1.8 µm | Agilent | 959757-902 |
| C18 guard column | 2.1 x 5 mm, 1.8 µm | Agilent | 821725-201 |
| ESI tune mix solution | n/a | Agilent | G1969-85000 |

### Table S2. LC gradient settings

| Time (min) | % B |
|---|---|
| 0 | 10 |
| 0.5 | 10 |
| 2 | 30 |
| 14 | 95 |
| 16.5 (stop time) | 100 |
| 6 (post time) | 10 |

### Table S3. Electrospray ionization source settings

| Parameter | Value | Units |
|---|---|---|
| Gas temperature | 230 | °C |
| Drying gas flow | 11 | L/min |
| Nebulizer gas pressure | 45 | psig |
| Sheath gas temperature | 350 | °C |
| Sheath gas flow | 11 | L/min |
| Vcap | 3500 | V |
| Nozzle voltage | 500 | V |

### Table S4. IMS-MS settings

| Parameter | Value | Units |
|---|---|---|
| Mass range | 50-1700 | *m/z* |
| Trap fill time | 3900 | µs |
| Trap release time | 300 | µs |
| Frame rate | 1 | frames/sec |
| IM transient rate | 17 | transients/frame |
| Max drift time | 60 | ms |
| TOF transient rate | 496 | transients/IM transient |
| Multiplexing pulse sequence length | 4 | bit |
| Drift tube entrance voltage | -1574 | V |
| Drift tube exit voltage | -224 | V |
| Rear funnel entrance voltage | -217.5 | V |
| Rear funnel exit voltage | -45 | V |

**Absolute and semi-quantitation methods.**

In the initial sample runs in 2022, concentrations of PFAS identified to Level 1a were calculated semi-quantitatively using single point internal calibration. In this single point concentration calculation, the relative abundance (light-to-heavy peak area ratio per g tissue) of an analyte is multiplied by the known concentration of a spiked internal standard to get the concentration of the native molecule. However, a key limitation of semi-quantitation is the inability to assess error over a range of concentrations. As increasing regulatory attention has been given to PFAS since this study was initiated in 2020, the need for accurate quantification has increased. Thus, absolute quantification was performed on the sample extracts in a new experiment in March 2024, after the initial non-targeted analyses were completed. To ensure that the sample extracts had not degraded in storage following the initial run, the reproducibility of relative abundances in all samples was compared to those of the original sample run (**Table ST4, Figure S1**). All samples were within 30% of the original values for PFOS, which is within the expected range of variation of our instrument.

Good accuracy and precision were obtained for the PFOS in the SRM samples as well. However, the other 3 PFAS with reference values available (PFNA, PFDA, and PFUdA) were below method LODs (**Table S5**). Because we could not use the reference material to validate our calibration curve accuracy for those PFAS, we opted to report PFOS values only because 1) we have high confidence in the accuracy of these concentrations and 2) PFOS is the most relevant PFAS to human exposure via fish consumption given that state consumption advisories exist are typically limited to PFOS only. Future NTA studies relevant to human PFAS exposure should include absolute quantification of the most relevant PFAS to continue increasing our understanding of human PFAS exposure routes.

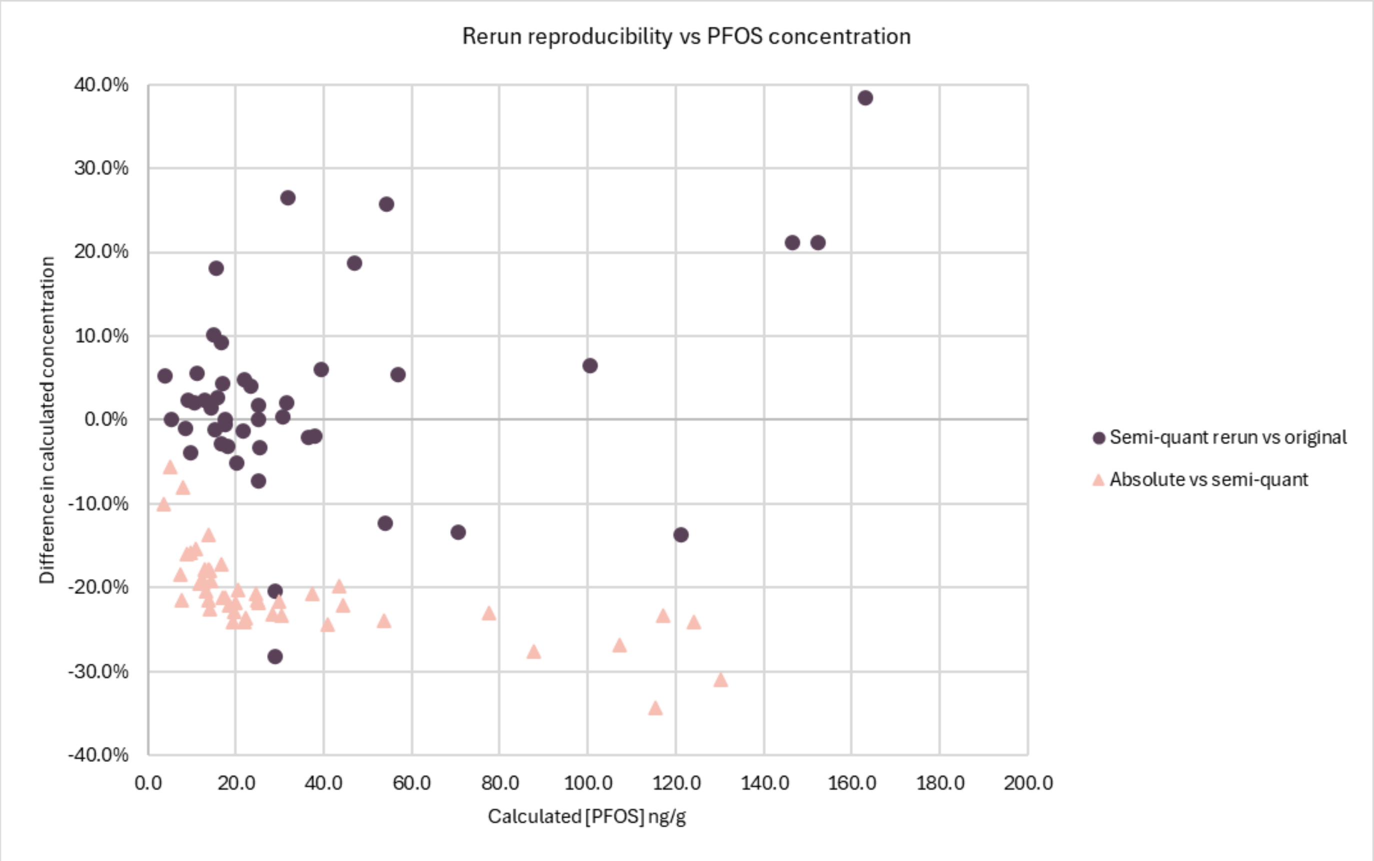

**Figure S1. Reproducibility of PFOS calculations.** The dark purple circles show the difference between the March 2024 rerun and the 2022 original run of the fish samples, comparing the PFOS concentrations calculated semi-quantitatively. Most of the rerun samples were within ± 30% of the original value, which is within the expected error of our instrument. The pink triangles show the difference between the PFOS concentrations calculated using the calibration curve and those calculated semi-quantitatively. Absolute quantification underestimated semi-quantitation in all cases, though the differences were less than 30% for almost all samples.

### Table S5. Accuracy and precision of PFAS quantitation

| | PFNA | PFDA | PFUdA | PFOS |
|---|---|---|---|---|
| **Limit of detection (ng/ml)** | 0.0499 | 0.0572 | 0.0592 | 0.0352 |
| **Reference value (ng/ml)** | 0.0182 | 0.0236 | 0.0254 | 0.5359 ± 0.0973 |
| **Accuracy[1]** | 235.1 to 290.7% | 237 to 266.4% | 233.2 to 257.9% | 101% |

[1] $Accuracy = \frac{Average\ measured\ concentration\ (ppb)}{NIST-reported\ concentration\ (ppb)} \times 100\%$, acceptance criteria 70-130%

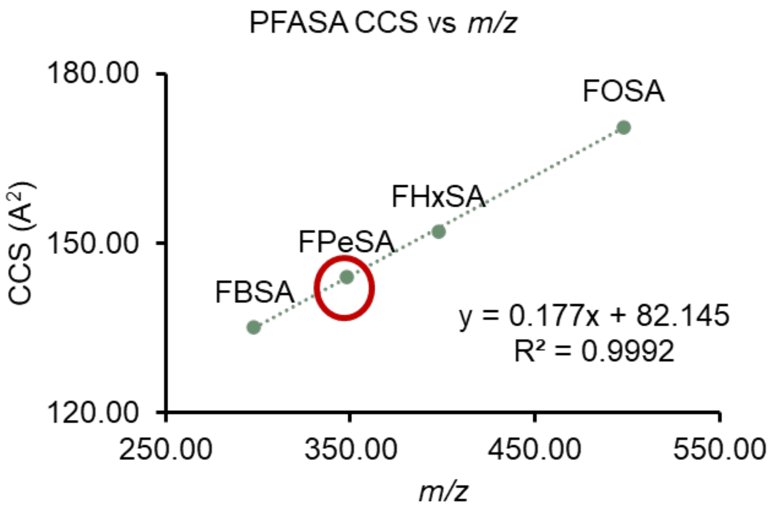

**Figure S2. Example of using CCS vs *m/z* trend lines to increase annotation confidence with PFAS.** We had previously obtained CCS values using FBSA, FHxSA, and FOSA standards. A standard is not available for FPeSA. A feature that matched the exact mass for FPeSA was detected in the NIST SRM 1947 samples, and the calculated CCS for this feature falls perfectly along the trendline expected for the perfluorosulfonamides.

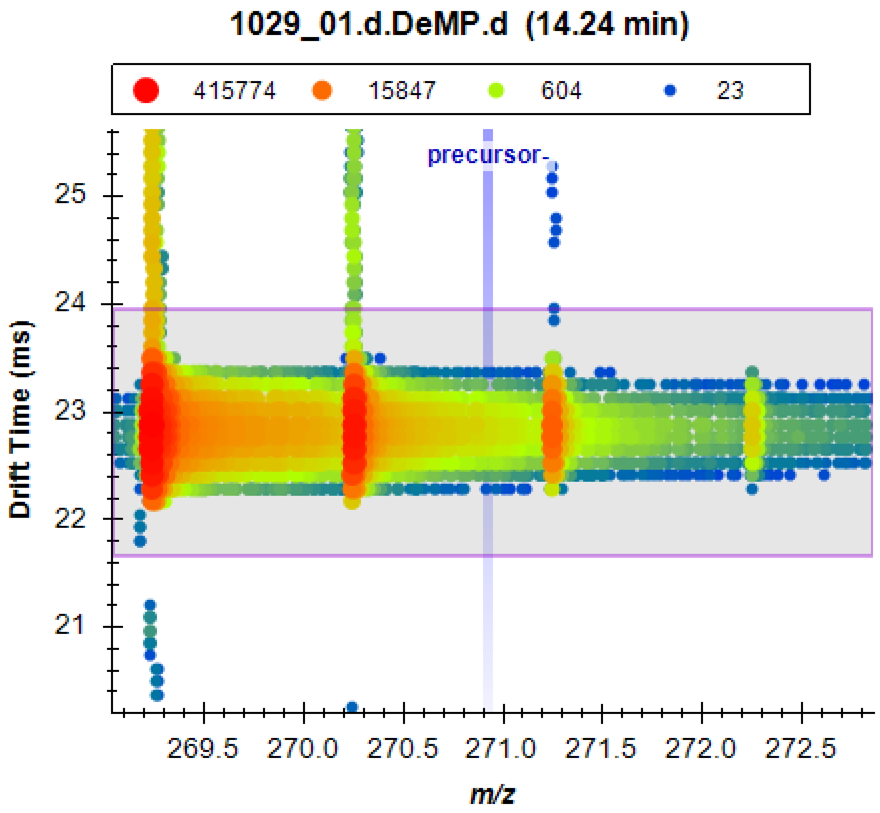

**Figure S3. Example of feature meeting criteria of possible PFAS but ultimately ruled out.** This feature (ID# 7855) at *m/z* 270.9264, RT 14.2 min, DT 22.8 ms was found using MassProfiler and met the filtering criteria we set in Fluoromatch-IM of negative mass defect, negative Kendrick mass defect, presence in a $CF_2$(n) homologous series, and abundance in fish samples ≥ 2.0 times the abundance in method blanks. The nested spectrum from one fish (sample 1029) shows *m/z* 270.9264 with a resolving power window of 10,000 in the vertical blue box, and the drift time 22.8 ms with a resolving power window of 20 in the horizontal pink box. The integrated peak area is the overlap of the *m/z* and drift time boxes. Clearly, the signal being integrated is tailing from the peak at *m/z* 270.25, and not likely to be a PFAS.

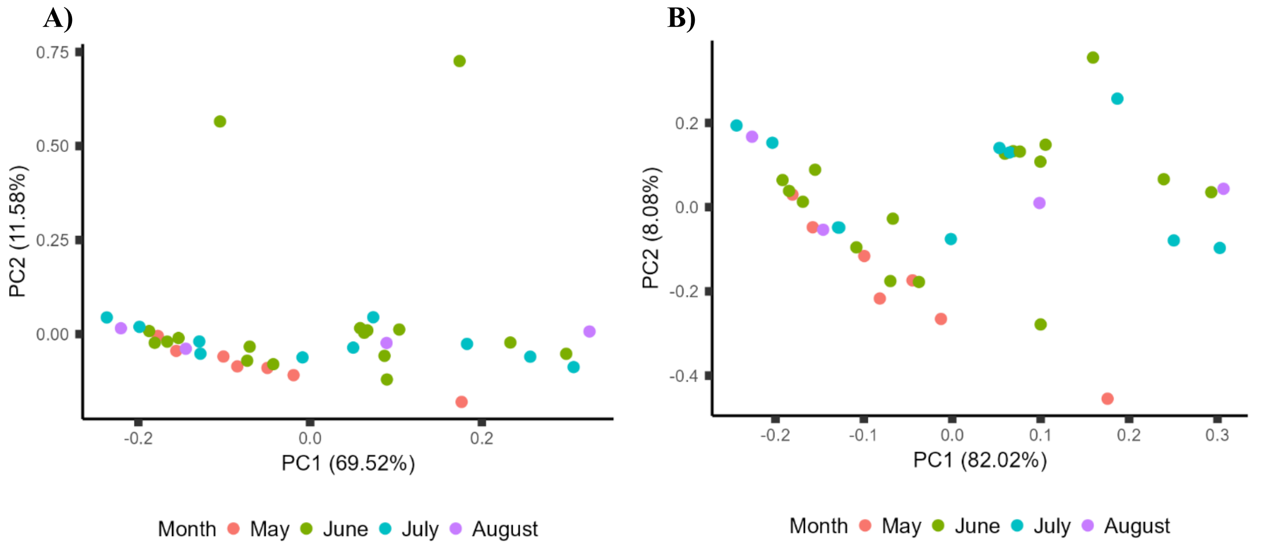

**Figure S4. Principle Component Analysis (PCA) of sampling date for *Lepomis* spp. fillet samples**, with each month shown in a different color. **A)** Evaluation based on all detected analytes. **B)** Evaluation based on the 10 analytes detected in more than 70% of fillets. No distinct separation by sampling month was observed. MANOVA as described in the main text confirmed no significant differences when comparing relative abundances of the 7 analytes detected in > 90% of fish (df = 3, Pillai's trace = 0.668, approx. F = 1.227, p = 0.249).

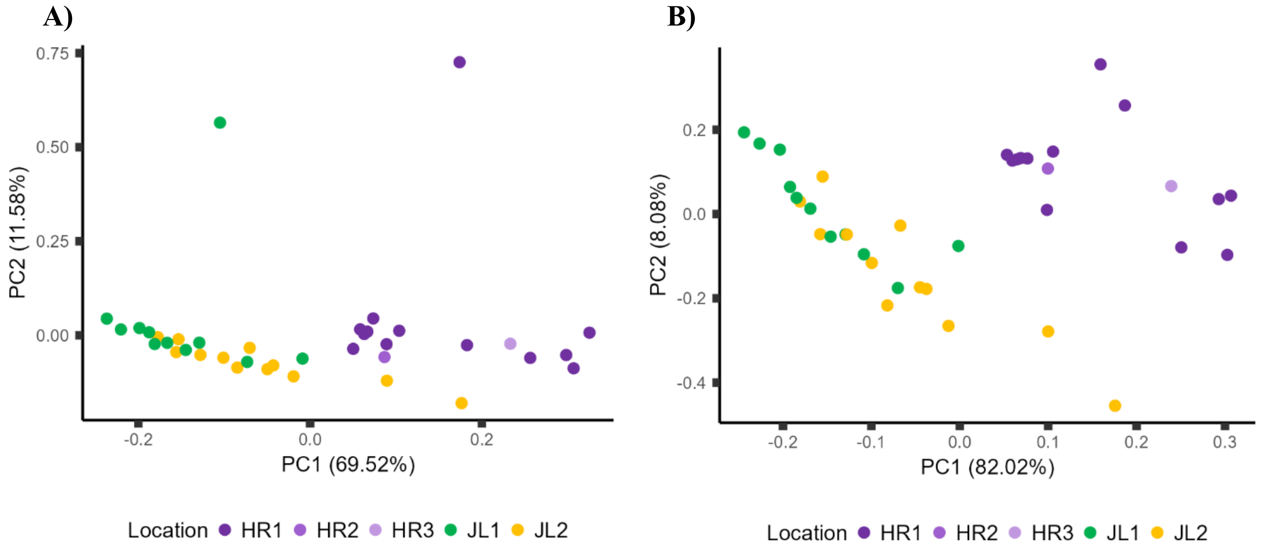

**Figure S5. PCA of *Lepomis* spp. sampling location**. **A)** Evaluation based on all detected analytes. **B)** Evaluation based on the 10 analytes detected in more than 70% of fillets. In both cases, the HR2 and HR3 samples clustered with the HR1 samples, and the HR samples separated from the JL samples. Distinct separation between the two JL sites was less clear.

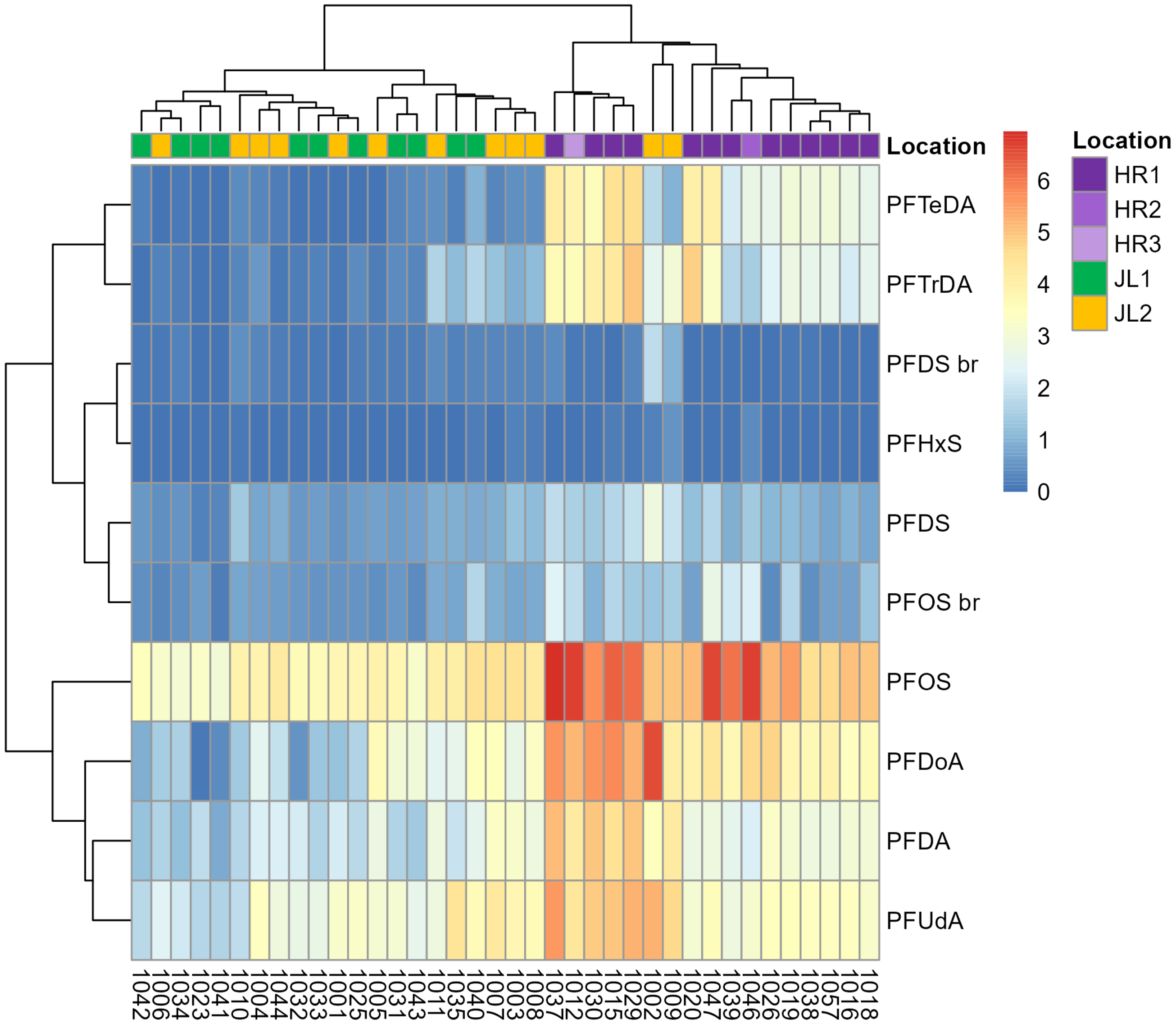

**Figure S6. Heat map of *Lepomis* spp. relative PFAS abundance at the 5 sampling sites**, using the 10 analytes detected in more than 70% of fillets. The HR2 and HR3 samples cluster tightly with the HR1 samples. The two JL2 samples that clustered the closest to the HR samples had particularly high PFOS and PFDoA abundance. Similarly to the PCA shown in Figure S3B, distinct separation between JL1 and JL2 was not observed.

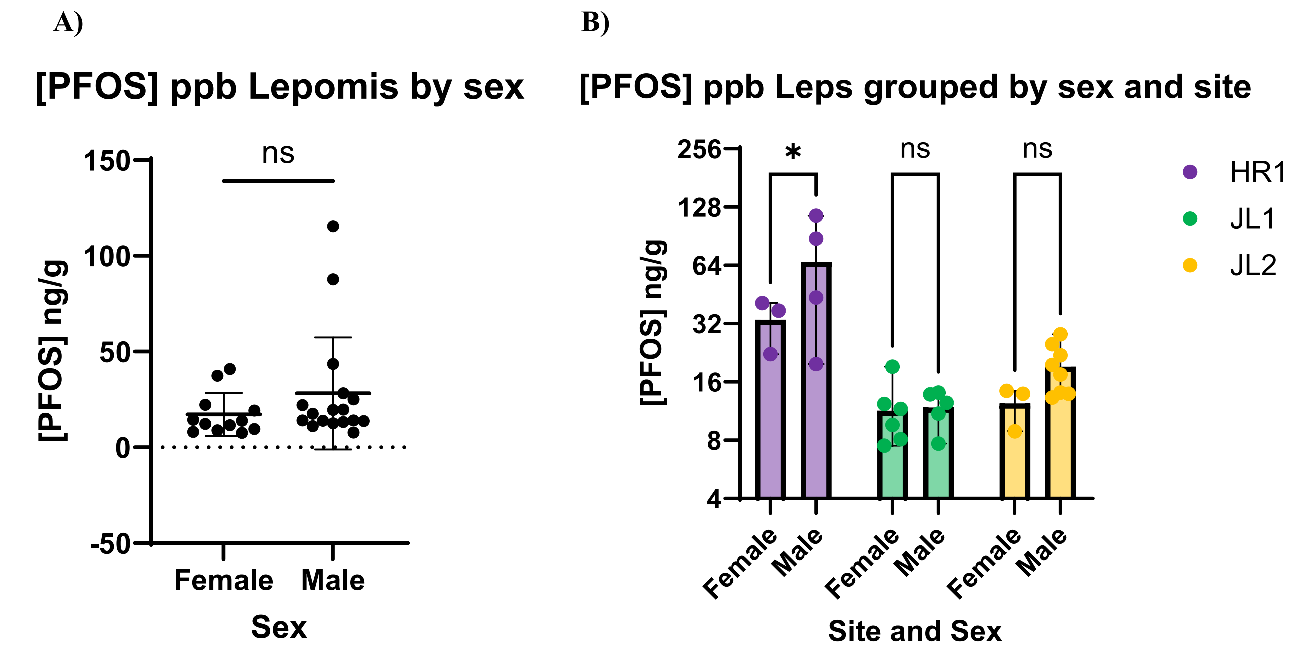

**Figure S7. Sex differences in PFOS concentration in *Lepomis* spp. sunfish fillets. A)** Student's t-test comparisons showed no significant differences by sex when all samples were combined ($p$ = 0.2265). **B)** Two-way ANOVA with Šídák's multiple comparisons test indicated a significantly higher concentration in males compared to females at HR1 (adjusted $p$ = 0.0411), but not at JL1 or JL2. However, given the small sample size and large variation in the samples, we have opted to group females and males together for all other analyses.

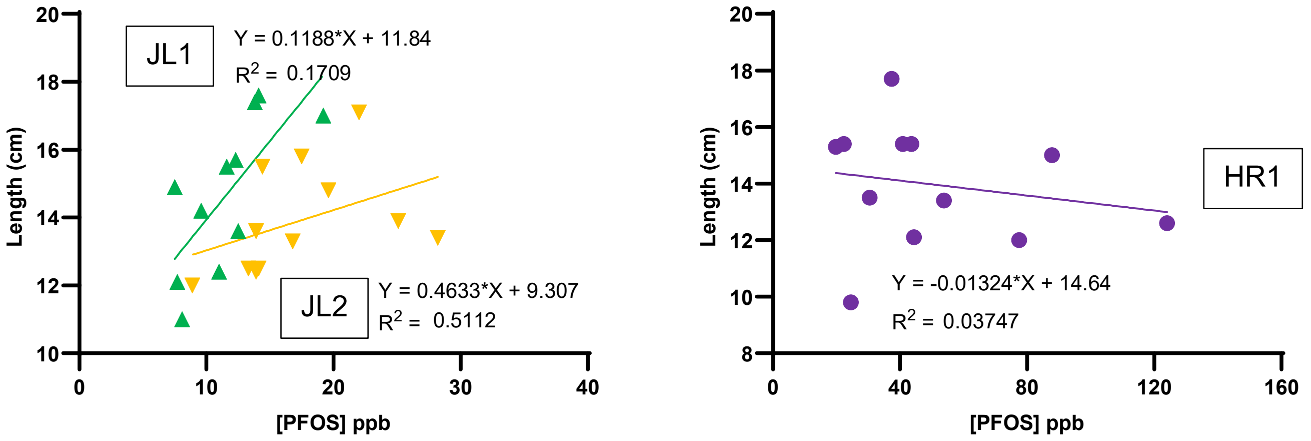

**Figure S8. Linear regression analysis of fish length and PFOS concentration in *Lepomis* spp. sunfish fillets.** Regression analysis indicated a significant correlation of fish length and PFOS concentration in fish fillets at site JL1 (p = 0.0134), but not at JL2 (p = 0.1816) or HR1 (p = 0.5466). Fish length is an indicator of fish age.
